# Contrary neuronal recalibration in different multisensory cortical areas

**DOI:** 10.1101/2022.09.26.509476

**Authors:** Fu Zeng, Adam Zaidel, Aihua Chen

**Affiliations:** Key Laboratory of Brain Functional Genomics (Ministry of Education), East China Normal University, 3663 Zhongshan Road N., Shanghai 200062, China; Gonda Multidisciplinary Brain Research Center, Bar-Ilan University, Ramat Gan, 5290002, Israel

**Keywords:** cross-modal, plasticity, self-motion, vestibular, visual, adaptation

## Abstract

The adult brain demonstrates remarkable multisensory plasticity by dynamically recalibrating information from multiple sensory sources. When a systematic visual-vestibular heading offset is experienced, the unisensory perceptual estimates recalibrate toward each other (in opposite directions) to reduce the conflict. The neural substrate of this recalibration is unknown. Here, we recorded single-neuron activity from the dorsal medial superior temporal (MSTd), parieto-insular vestibular cortex (PIVC), and ventral intraparietal (VIP) areas in three male rhesus macaques during visual-vestibular recalibration. Both visual and vestibular tuning in MSTd recalibrated-each according to their respective cues’ perceptual shifts. Vestibular tuning in PIVC also recalibrated together with corresponding perceptual shifts (cells were not visually tuned). By contrast, VIP neurons demonstrated a unique phenomenon: both vestibular and visual tuning recalibrated according to vestibular perceptual shifts. Such that, visual tuning shifted, surprisingly, contrary to visual perceptual shifts. Therefore, while unsupervised recalibration (to reduce cue conflict) occurs in early multisensory cortices, higher-level VIP reflects only a global shift, in vestibular space.

**In Brief:** The neural bases of multisensory plasticity are currently unknown. Here, Zeng et al. studied neuronal recalibration to a systematic visual-vestibular cue conflict. In multisensory cortical areas MSTd and PIVC, single-unit responses to visual and vestibular stimuli recalibrated to reduce the cue conflict, along with their respective unisensory perceptual shifts. By contrast, in higher-level VIP, both visual and vestibular neuronal responses recalibrated with vestibular perceptual shifts. This led to a surprising recalibration of visual responses opposite in direction to visual perceptual shifts. This exposes differential aspects of multisensory plasticity across multisensory cortical areas, and reveals a novel hybrid of visual responses within a vestibular reference frame in parietal neurons.

**Highlights:**

- In the presence of a systematic heading conflict, visual and vestibular cues recalibrate towards one another to reduce the conflict.
- In MSTd, neuronal responses to vestibular and visual cues recalibrated, each according to their respective cues’ perceptual shifts.
- In PIVC, vestibular responses recalibrated according to vestibular perceptual shifts (cells were not visually tuned).
- In VIP, neuronal responses to both vestibular and visual cues recalibrated together with vestibular perceptual shifts (opposite in direction to visual perceptual shifts).
- Profound differences in neuronal recalibration expose different functions across multisensory cortical areas.

## Introduction

Our different sensory systems each continuously adapt to changes in the environment (Webster, 2012). Thus, to maintain stable and coherent perception in a multisensory and ever-changing world, the brain needs to dynamically adjust for sensory discrepancies between the different modalities. This process of multisensory recalibration takes place continually, and is perhaps more fundamental than multisensory integration because integration would not be beneficial when the underlying cues are biased. While the neural bases of multisensory integration have received a lot of attention (Chen et al., 2013a; Ernst and Banks, 2002; Ernst and Bülthoff, 2004; Ernst and Di Luca, 2011; Gu et al., 2008; Stein et al., 2014), the neural bases of multisensory recalibration have been explored to a much lesser degree.

Cross-modal recalibration has been observed in a variety of multisensory settings. One well-known example is the ventriloquist aftereffect (VAE), in which exposure to a consistent spatial discrepancy between auditory and visual stimuli induces a subsequent shift in the perceived location of sounds (Bertelson and De Gelder, 2004; Canon, 1970; Kramer et al., 2020; Radeau and Bertelson, 1974; Recanzone, 1998; Watson et al., 2021). Also, the rubber-hand illusion (RHI) leads to an offset in hand proprioception in the direction of the visually observed rubber hand (Abdulkarim et al., 2021; Botvinick and Cohen, 1998; Kennett et al., 2001; Thériault et al., 2022; Tsakiris and Haggard, 2005). Although it was initially thought that only the non-visual cues recalibrate to vision (visual dominance; (Brainard and Knudsen, 1993; Rock and Victor, 1964), further work in a variety of paradigms has revealed both visual and non-visual recalibration (Atkins et al., 2003; Burge et al., 2010; Lewald, 2002; van Beers et al., 2002; Zaidel et al., 2011).

Most of what we know about multisensory recalibration is described at the behavioral level (Burge et al., 2008; Burge *et al*., 2010; Lewald, 2002), with little known about its neuronal underpinnings. Recent EEG (Park and Kayser, 2021) and fMRI (Zierul et al., 2017) studies in humans have shed some light on this question. However, these methods lack the resolution to probe recalibration at the level of single neurons. A series of classic studies by Eric Knudsen and colleagues investigated multisensory plasticity at the neuronal and circuit levels, in the barn owl (Knudsen, 2002; Knudsen and Brainard, 1991; Linkenhoker and Knudsen, 2002). They found profound neuronal plasticity in juvenile owls reared with prismatic lenses that systematically displaced their field of view. In that case, the auditory space map in the optic tectum was recalibrated to be aligned with the displaced visual field (Knudsen and Brainard, 1991). However, multisensory plasticity is not limited to the development, and the neuronal bases of how multiple sensory systems continuously adapt to one another in the adult brain remain fundamentally missing.

Self-motion perception (the subjective feeling of moving through space) relies primarily on visual and vestibular cues (Butler et al., 2015; Butler et al., 2010; de Winkel et al., 2010; Fetsch et al., 2012; Fetsch et al., 2009; Gu et al., 2007; Warren et al., 1988). Multisensory integration of visual and vestibular signals can improve heading perception (Burge *et al*., 2010; Butler *et al*., 2015; Dokka et al., 2015; Gu *et al*., 2008). However, conflicting or inconsistent visual and vestibular information often leads to motion sickness (Oman, 1990; Reason and Brand, 1975). Interestingly, this subsides after prolonged exposure to the sensory motion conflict, presumably through brain mechanisms of multisensory recalibration (Held, 1961; Shupak and Gordon, 2006). Thus, self-motion perception – a vital skill for everyday function with intrinsic plasticity – offers a prime substrate to study cross-sensory recalibration.

We previously investigated and found robust (behavioral) recalibration of both visual and vestibular cues in response to a systematic vestibular-visual heading discrepancy (Zaidel *et al*., 2011). In that paradigm, no external feedback was given. Thus, the need for recalibration arose solely because of the cue discrepancy (we therefore call this condition *unsupervised*). The subjects (humans and monkeys) recalibrated both visual and vestibular perceptual estimates by shifting them toward each other, to reduce the conflict. This is in line with the notion that unsupervised recalibration aims to maintain “internal consistency” between the cues (Burge *et al*., 2010). However, the neuronal basis of this everyday multisensory plasticity is unknown.

In a complementary behavioral study, we tested *supervised* self-motion recalibration, by providing external feedback regarding cue accuracy (Zaidel et al., 2013). There we found that supervised recalibration is a high-level cognitive process that compares the combined-cue (multisensory) estimate to feedback from the environment. This resulted in ‘yoked’ recalibration of both cues, in the same direction, to reduce conflict between the combined estimate and external feedback. We subsequently also investigated the neuronal substrate of *supervised* recalibration (Zaidel et al., 2021). We found robust recalibration of both vestibular and visual neuronal tuning in the monkey ventral intraparietal (VIP) cortex, such that tuning for both cues shifted together, in accordance with the behavior. However, because in that paradigm both cues recalibrate in the same direction (yoking), neuronal tuning was also expected to shift in the same direction for both cues. Thus, differential aspects of neuronal recalibration for the individual cues could go undetected.

By contrast, in unsupervised recalibration, vestibular and visual cues shift in opposite directions (Zaidel *et al*., 2011). Therefore, the unsupervised paradigm can better expose differences in the way that individual cues recalibrate to one-another in the brain. Because unsupervised recalibration occurs in the absence of external feedback, it is presumed to reflect implicit changes in perception. Thus, we expected to see its effects relatively early in the vestibular-visual integration hierarchy, and that these effects would propagate to higher-level areas. Unsupervised recalibration of single neurons in single behavioral sessions, has not been tested before. The resulting psychometric shifts are smaller (vs. supervised recalibration). Thus detecting its neuronal correlates is challenging, but imperative, to understand the neural bases of adult cross-sensory plasticity. Thus, the aim of this study was to test unsupervised recalibration of visual and vestibular neuronal tuning, and how it may differ across multisensory cortical areas.

Two relatively early multisensory cortical areas involved in self-motion perception are the medial superior temporal area (MSTd) and the parietal insular vestibular cortex (PIVC). Neurons in MSTd respond to large optic flow stimuli, conducive to the visual perception of self-motion (Gu et al., 2006). Vestibular responses are also present in MSTd, however visual self-motion signals dominate (Gu *et al*., 2008; Gu et al., 2012). PIVC has strong vestibular responses, without strong tuning to visual optic flow (Chen et al., 2010). Therefore, we expected to see perceptual shifts resulting from unsupervised calibration in MSTd and PIVC. Area VIP also has robust responses to visual and vestibular self-motion stimuli, however, it is marked by strong choice signals (Chen et al., 2016; Gu, 2018; Zaidel et al., 2017). It is thus considered a higher-level multisensory area involved in additional (currently not fully understood) cognitive functions. Different types of multisensory recalibration observed in these different multisensory areas can provide important insights into their differential underlying functions. Thus, in this study, we focused on these three multisensory cortical areas. We examined whether and how their visual and vestibular neural tuning changed in accordance with corresponding behavioral shifts during a single session (∼1hr) of unsupervised cross-sensory recalibration.

## Results

Three monkeys performed a task of heading discrimination in a paradigm that elicits unsupervised cross-sensory (vestibular-visual) recalibration. Simultaneous to behavioral performance, we recorded from single neurons extracellularly in areas MSTd (upper bank of the superior temporal sulcus, n=83: 19 from monkey D, 64 from monkey K), PIVC (upper bank and the tip of the lateral sulcus, n=160: 91 from monkey D, 69 from monkey B), and VIP (lower bank and tip of the intraparietal sulcus, n=118: 103 from monkey D, 15 from monkey B). The paradigm followed the same methodology as our previous behavioral study (Zaidel *et al*., 2011). It consisted of three consecutive blocks: pre-recalibration (**Fig. 1A**), recalibration (**Fig. 1B**), and post-recalibration (**Fig. 1C**). We first (in the next section) present the monkeys’ perceptual recalibration results. Thereafter, we present the neural correlates thereof.

**Figure 1.**
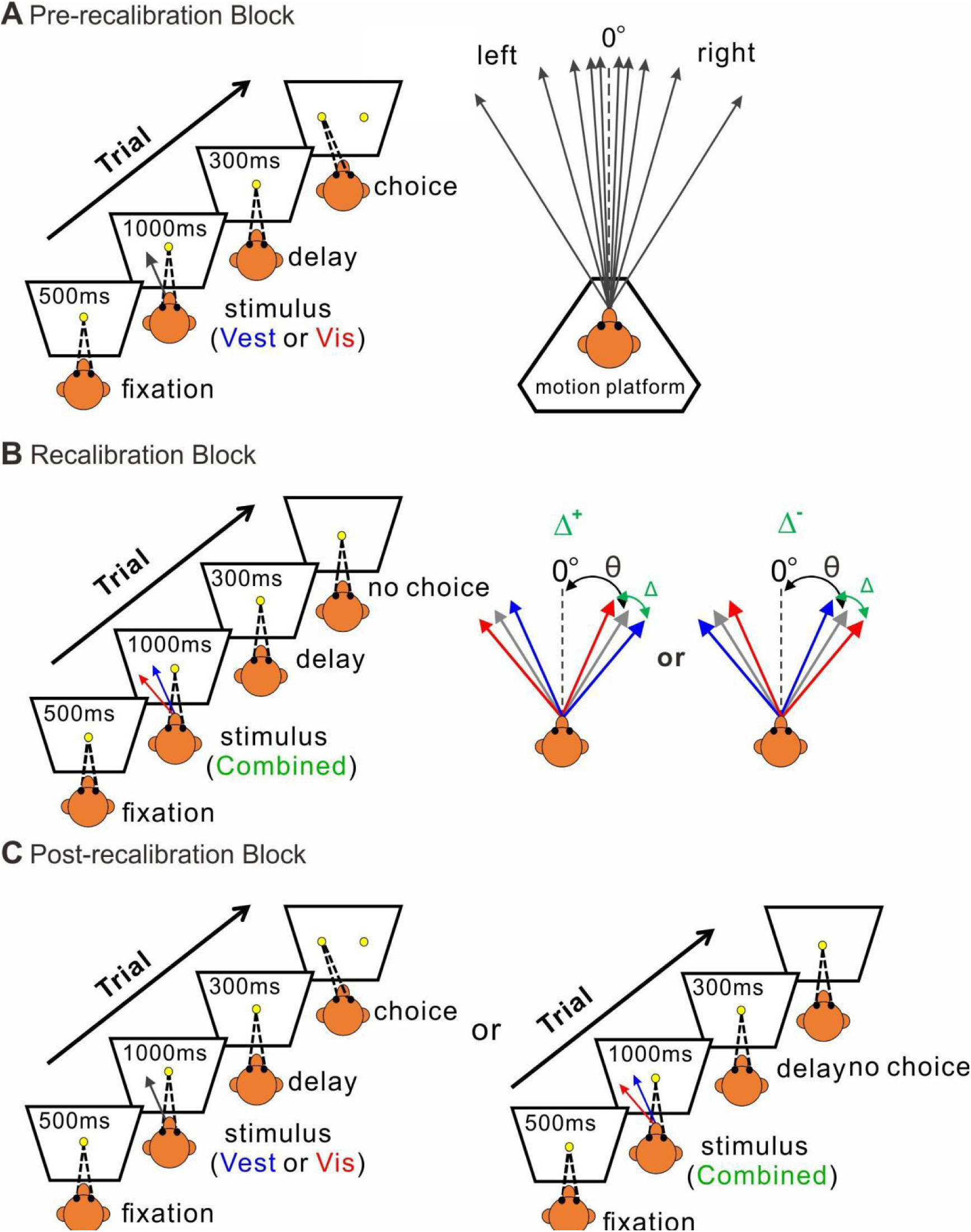
Multisensory recalibration paradigm. **(A)** Pre-recalibration block. The vestibular stimulus was provided by the motion platform (schematic on the right), and the visual stimulus was optic-flow simulation of self-motion (without motion of the platform) presented on a screen in front of the monkey (schematic on the left). The self-motion stimuli comprised linear motions (of either vestibular or visual stimuli) in a primarily forward direction, with slight deviations to the right or left (black arrows, schematic on the right). Monkeys were required to fixate on a central target (yellow circle) presented on the screen during the stimulus and then to report their perceived heading by making a saccade to one of two choice targets (left or right relative to straight ahead). **(B)** Recalibration block. Vestibular and visual stimuli were presented together (“combined”) with a systematic discrepancy (Δ) between the vestibular and visual headings. The blue and red arrows represent the vestibular and visual headings, respectively. The gray arrows represent the combined cue headings (in between the vestibular and visual cues) and the black dashed lines represent straight ahead. **(C)** Post-recalibration block. The single-cue trials (like in A) were interleaved with combined-cue trials (like in B).

### Both vestibular and visual cues recalibrate toward each other

**Figure 2** shows example psychophysical data from two experimental sessions. Replicating our previous behavioral results (Zaidel *et al*., 2011), we found that both visual and vestibular psychometric functions shifted in the direction required to reduce the cue conflict. Namely, when the vestibular and visual heading stimuli were systematically offset, such that they consistently deviated to the right and the left, respectively (Δ^+^, **Fig. 2A**), the vestibular post-recalibration curve (blue) was shifted rightward vs. pre-recalibration (black). Note that a rightward shift of the psychometric curve indicates a *leftward* perceptual shift (identified by a lower propensity for ‘rightward’ choices at 0° heading for the blue curve). Complementarily, the visual post-recalibration psychometric curve (red) shifted leftward vs. pre-recalibration (black), albeit to a lesser degree, indicating a *rightward* perceptual shift. In a reverse manner, when the vestibular and visual heading stimuli were offset to the left and right respectively (Δ^−^, **Fig. 2B**), the vestibular post-recalibration curve (blue) shifted to the left, and the visual post-recalibration curve shifted to the right.

**Figure 2.**
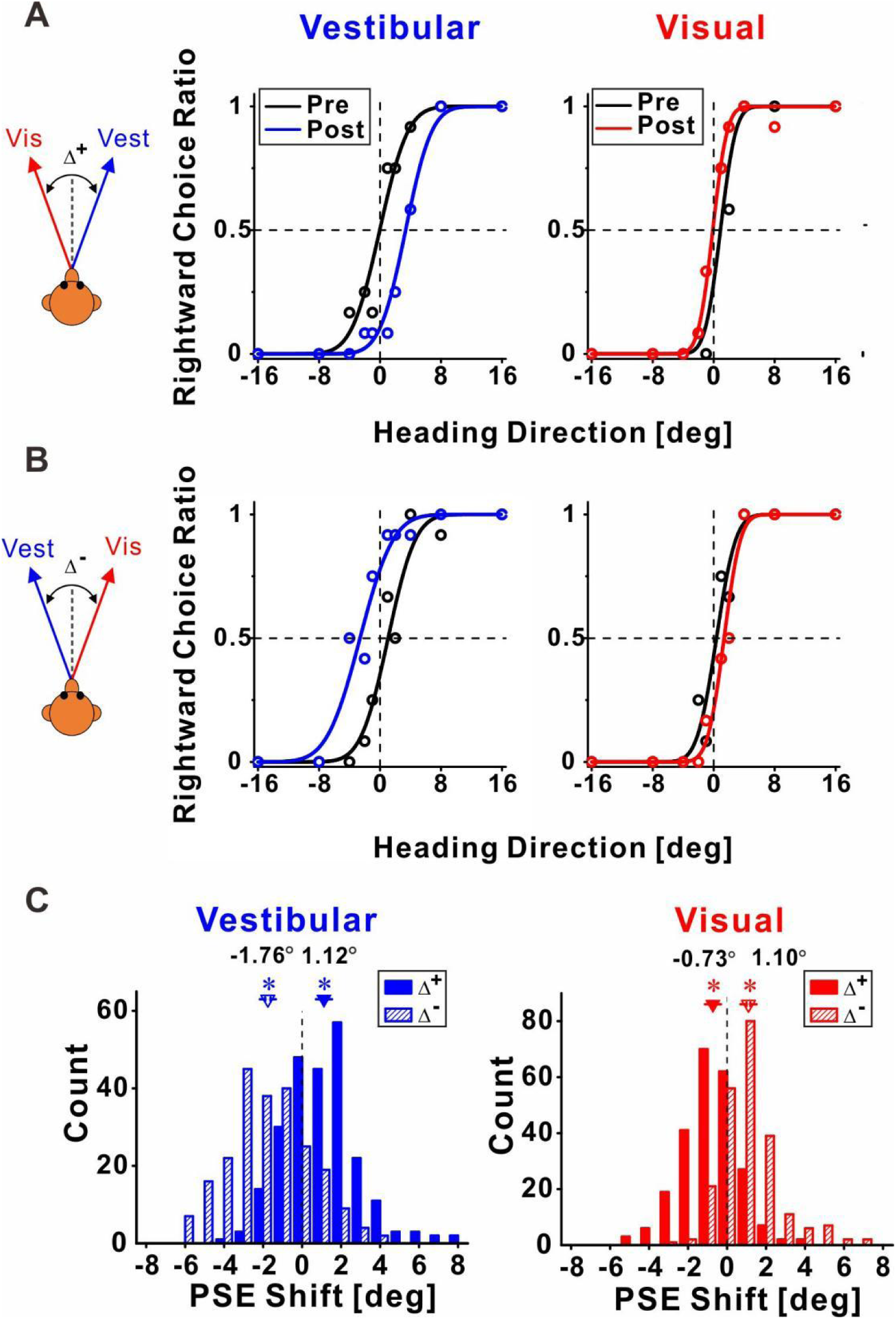
Multisensory recalibration behavior. **(A, B)** Example psychometric plots represent the ratio of the monkeys’ rightward choices, as a function of stimulus heading direction. Data (circles) were fitted with cumulative Gaussian functions (solid lines). Pre-recalibration performance is presented for vestibular and visual cues (in the left and right columns, respectively) by the black curves. After recalibration, vestibular and visual cues were shifted (blue and red curves, respectively). Behavior for positive and negative delta (Δ + and Δ -, respectively) are presented in A and B, respectively. (C) Blue and red histograms represent the vestibular and visual PSE shift distributions, respectively. Solid and slash-textured histograms indicate positive and negative Δ, respectively. Inverted triangles (▾) and error bars represent mean SEM shifts. The numbers above triangles are the mean PSE shift. Asterisk symbols indicate significant shifts (p < 0.05). In vestibular cue, p = 2.54 × 10^−15^ for Δ+ condition, and p = 1.77 × 10^−29^ for Δ- condition, respectively. In visual cue, p = 1.60 × 10^−10^, n = 241 sessions for Δ+ condition, and p = 2.94 × 10^−21^, n = 227 sessions for Δ- condition, respectively, paired t-test.

These behavioral shifts were quantified by the difference between the post- vs. pre-recalibration curves’ PSEs (points of subjective equality). Each psychometric curve’s PSE was detected by the heading at which it crosses y = 0.5 (marked by horizontal dashed lines in **Fig. 2**). The vestibular and visual psychometric shifts were: 3.40° and −1.01°, respectively in **Figure 2A**, and −3.68° and 1.00°, respectively, in **Figure 2B**. Thus, in both cases (**Fig. 2A, B**), both the vestibular and the visual cues shifted in the direction required to reduce the cue conflict (i.e. in opposite directions). Also, the vestibular shifts were larger (vs. visual).

These findings were consistent across sessions, as shown by the distributions of the vestibular and visual PSE shifts (solid bars for Δ^+^ and striped bars for Δ^−^) in **Figure 2C**. The vestibular PSEs were shifted significantly to the right for the Δ ^+^ condition (mean ± SE = 1.12° ± 0.12°; *p* = 2.54 × 10^−15^, paired t-test). And shifted significantly to the left for the Δ^−^ condition (mean ± SE = −1.76° ± 0.14°; *p* = 1.77 × 10^−29^, paired t-test). The visual PSEs were shifted significantly to the left for the Δ^+^ condition (mean ± SE = −0.73° ± 0.11°; *p* = 1.60 × 10^−10^, paired t-test). And shifted significantly to the right for the Δ^−^ condition (mean ± SE = 1.10° ± 0.10°; *p* = 2.94 × 10^−21^, paired t-test). Thus, consistent with our previous study, both cues shifted (in opposite directions) to reduce the cue conflict.

Comparing the vestibular vs. visual shift magnitudes (using absolute values, pooled across Δ^+^ and Δ^−^ conditions) demonstrated significantly larger vestibular vs. visual shifts (1.43° ± 0.09 and 0.91° ± 0.07°, respectively; *p* = 3.40 × 10^−5^, paired t-test). This result is also consistent with our previous study. Thus, the behavioral results from the original study (performed in the Angelaki laboratory) were replicated in these experiments (in the Chen laboratory) using a new set of monkeys, with simultaneous neuronal recording. In the following sections, we present how neuronal responses in areas MSTd, PIVC, and VIP (**Fig. 3**, **4,** and **5**, respectively) recalibrated in comparison to the behavioral shifts.

**Figure 3.**
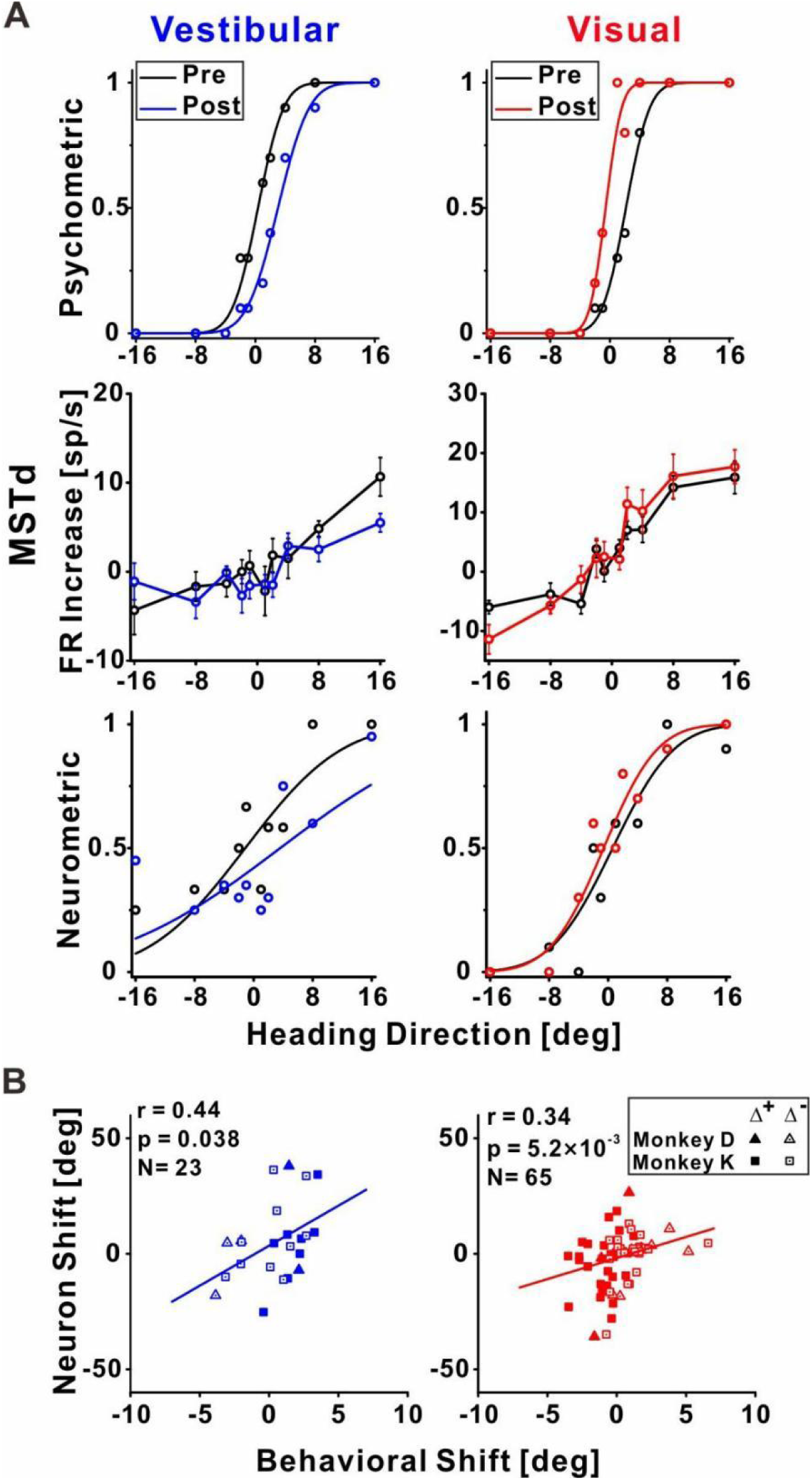
MSTd neuronal recalibration. **(A)** An example recalibration session (Δ^+^) with simultaneous recording from MSTd. The top row depicts the behavioral responses, pre-, and post-recalibration. The vestibular psychometric curve shifted 3.01° (to the right) and the visual curve shifted −2.71° (to the left). Neuronal responses (second row) as a function of heading (pre- and post-recalibration). Circles and error bars represent average firing rates (FRs, baseline subtracted) ± SEM. The third row shows corresponding neurometric functions with best-fitting cumulative Gaussian functions. The vestibular neuronal shift was 4.73° (to the right) and the visual neuronal shift was −1.22° (to the left). **(B)** Correlations between neuronal PSE shifts and behavioral PSE shifts for the vestibular and visual cues (left and right, respectively). Only cells with significant tuning (p < 0.05, Pearson correlation between firing rate and heading) in either pre- or post-recalibration blocks were included here. Solid symbols represent Δ+ and open symbols represent Δ-. The solid lines illustrate the regression lines of the data. p = 0.038, n = 23 neurons for vestibular cue, p = 5.2 × 10^−15^, n = 65 neurons for visual cue, Pearson correlation.

**Figure 4.**
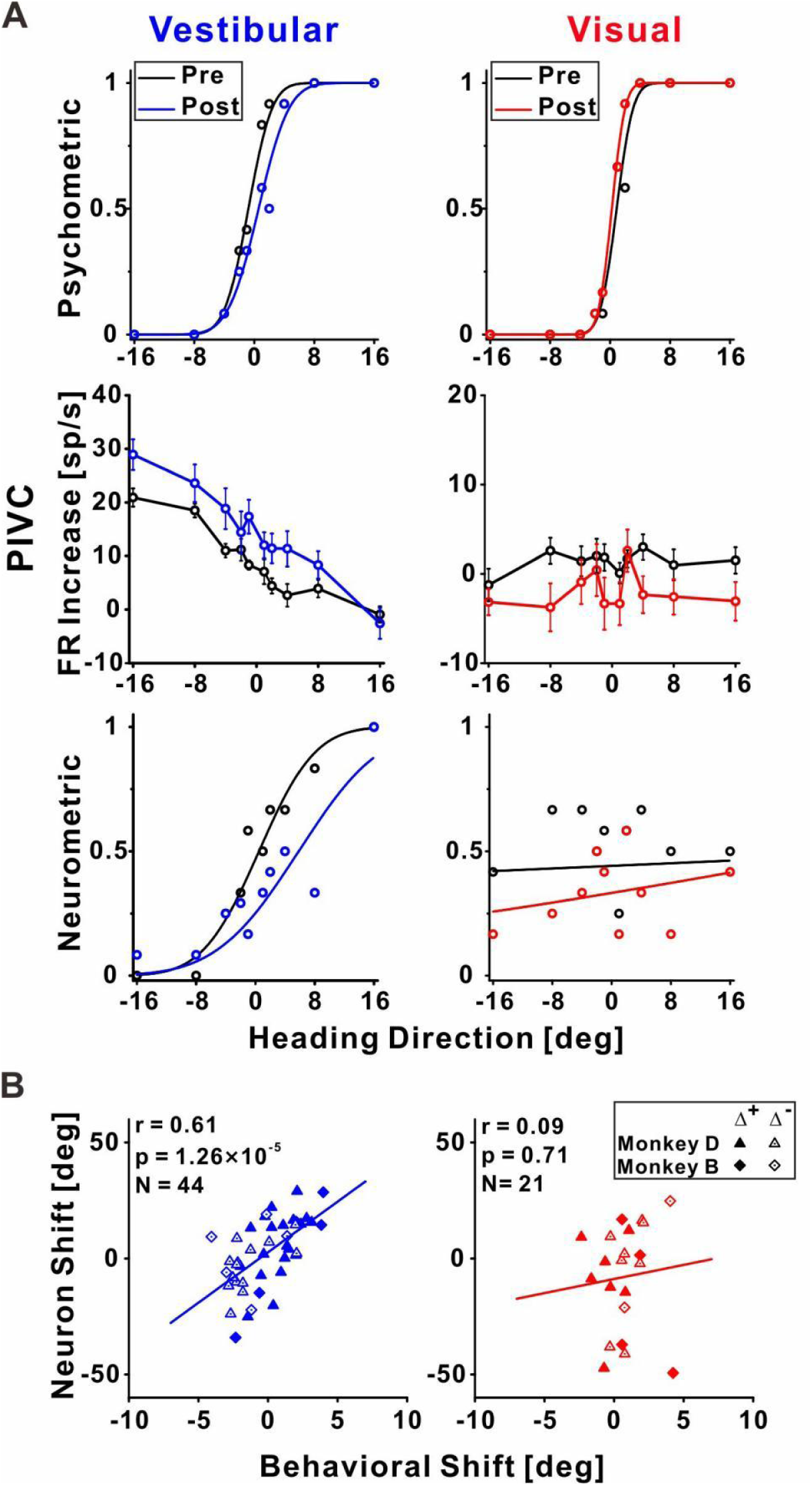
PIVC neuronal recalibration. **(A)** An example recalibration session (Δ^+^) with simultaneous recording from PIVC (conventions are the same as Figure 3). The vestibular and visual psychometric curves shifted 1.37 and −0.51 (to the right and left, respectively). The vestibular neurometric curve shifted 5.37 ° (to the right). **(B)** Correlations between neuronal PSE shifts and behavioral PSE shifts for the vestibular and visual cues. p = 1.26 × 10^−5^, n = 44 neurons for vestibular cue, p = 0.71, n = 21 neurons for visual cue, Pearson correlation.

**Figure 5.**
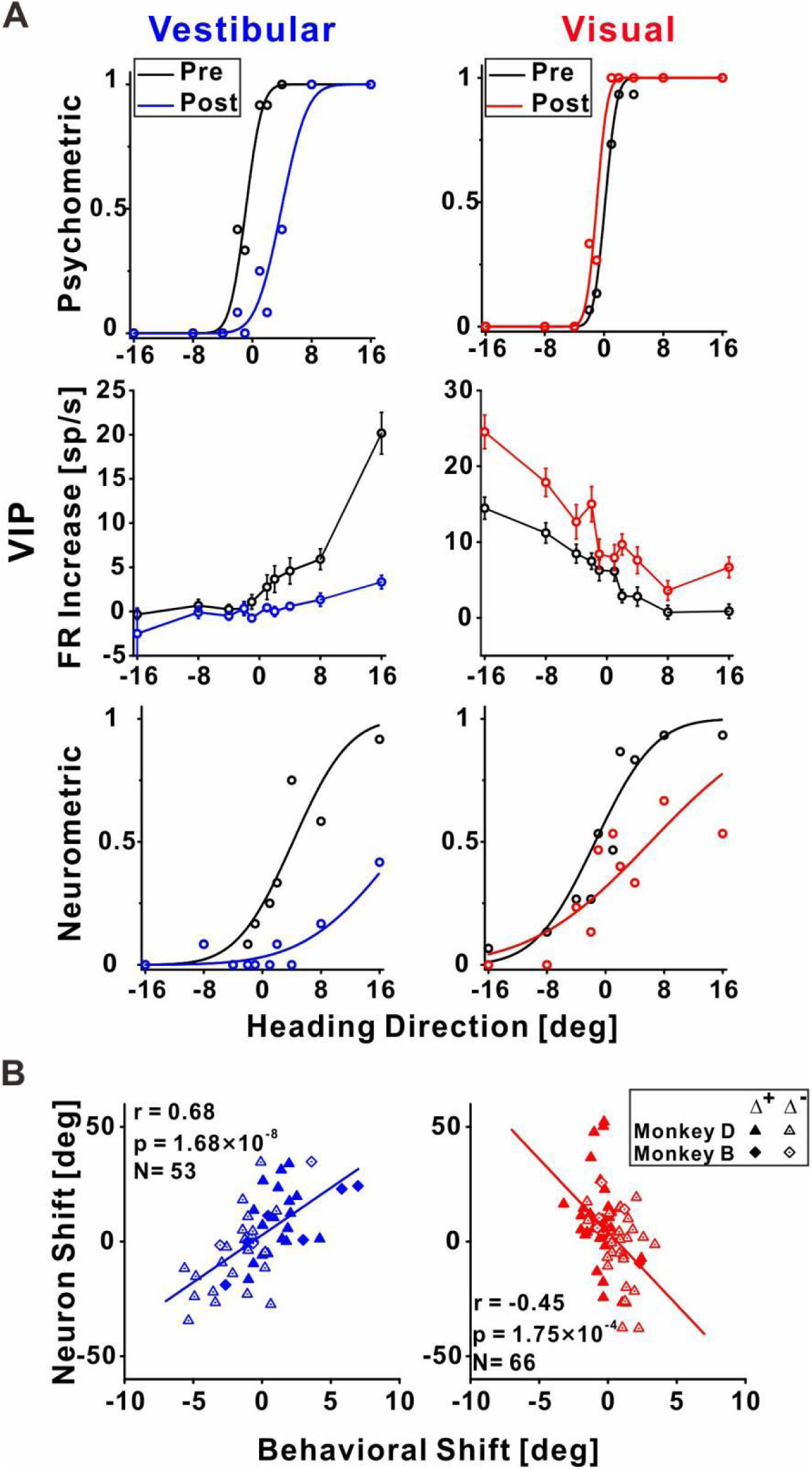
VIP neuronal recalibration. **(A)** An example recalibration session (Δ^+^) with simultaneous recording from VIP (conventions are the same as Figure 3). The vestibular and visual psychometric curves shifted 4.81 ° and −1.13 (to the right and left, respectively). The vestibular and visual neurometric curves shifted 15.18 ° and 7.58 °, respectively (both to the right). **(B)** Correlations between neuronal PSE shifts and behavioral PSE shifts for the vestibular and visual cues. p = 1.68 × 10^−8^, n = 53 neurons for vestibular cue, p = −0.45 × 10^−4^, n = 66 neurons for visual cue, Pearson correlation.

### Both vestibular and visual neuronal tuning in MSTd recalibrate with perceptual shifts

Responses of an example neuron recorded from MSTd during unsupervised recalibration are presented in **Figure 3A**. Behaviorally, the vestibular PSE shifted rightward and the visual PSE shifted leftward (upper panel, **Fig. 3A**). Shifts in neuronal tuning could be subtle, therefore we used neurometrics to expose and quantify the neuronal shifts. Specifically, we calculated neurometric responses for the heading stimuli using the neuron’s firing rates and ROC analysis, and fit these with a cumulative Gaussian function (for method details, see Gu et al., 2007). PSEs were then extracted, similar to the psychometric curves. Neurometric curves for this example neuron are presented in the third row of **Figure 3A**. For this MSTd neuron, the vestibular neurometric shifted to the right, while the visual neurometric shifted to the left. Thus, the shifts in vestibular and visual tuning were consistent with the behavioral shifts.

Across the population (**Fig. 3B**) MSTd neuronal shifts were significantly correlated with the behavioral shifts, both for vestibular and visual cues (*r* = 0.44, *p* = 0.038, N = 23, and *r* = 0.34, *p* = 5.2 × 10^−3^, N = 65, respectively; Pearson correlations). Therefore, in area MSTd neuronal recalibration occurs in accordance with perceptual recalibration, both for vestibular and visual cues.

### Vestibular neuronal tuning in PIVC recalibrates with perceptual shifts

In PIVC, a similar result was observed for vestibular tuning. The example vestibular neurometric curve **(Fig. 4A**, bottom left) shifted to the right, which was consistent with the behavioral shift **(Fig. 4A**, top left). Across the population of PIVC neurons, a significant positive correlation was seen between the neuronal and behavioral shifts for the vestibular cue (*r* = 0.61, *p* = 1.26 × 10^−5^, N = 44, Pearson correlation; **Fig. 4B**, left panel).

In terms of visual tuning, this example neuron (and the other PIVC neurons) did not demonstrate robust visual responses (**Fig. 4A**, middle right). However, we still applied the same neurometric analysis for visual responses, using PIVC neurons that passed the significance criterion for visual tuning (albeit weak). No significant correlation was seen between the neuronal and behavioral shifts for the visual cue (*r*=0.09, *p*=0.71, N=21, Pearson correlation). Thus, in PIVC, the primary cortical region involved in vestibular function, neuronal tuning shifts were consistent with perceptual shifts, for the vestibular cue.

### Vestibular and visual neuronal tuning in VIP both follow vestibular perceptual shifts

**Figure 5A** presents an example neuron from VIP. For the vestibular cue, the neuronal tuning curve shifted rightward (**Fig. 5A**, bottom left), in accordance with the vestibular behavioral shift (**Fig. 5A**, top left). Surprisingly, the visual neurometric curve also shifted rightward (**Fig. 5A**, bottom right). This was unexpected because the visual psychometric curve shifted leftward (**Fig. 5A**, top right). Thus, while the vestibular and visual behavioral psychometric curves shifted in opposite directions (toward each other) the vestibular and visual neurometric curves shifted together, in accordance with the vestibular (not visual) behavioral shift.

Across the population of VIP neurons, the vestibular neurometric shifts were significantly positively correlated with the vestibular behavioral shifts (*r* = 0.69, *p* = 1.68× 10^−8^, N = 53, Pearson correlation; **Fig. 5B**, left). Like in MSTd and PIVC, the positive correlation coefficient indicates that neuronal and behavioral curves shifted in the same direction for the vestibular cue. By contrast, the visual neurometrics in VIP shifted in the opposite direction to the visual behavioral shifts. At the population level neuronal and behavioral shifts for the visual cue were negatively correlated (*r* = −0.45, *p* = 1.75 × 10^−4^, Pearson correlation, N = 66; **Fig. 5B**, right). This exposes a striking mismatch between visual neuronal responses in VIP and visual perceptual function. It also exposes a striking mismatch between visual tuning in MSTd (which shifted in the same direction as visual perception) vs. VIP (which shifted contrary to visual perception).

To test whether this mismatch between behavior and tuning for visual cues in VIP relates to specific subtypes of neurons, we sorted the VIP data into three subsets: neurons with multisensory (vestibular and visual) responses, and two groups with unisensory (only vestibular or only visual) responses (**Supplemental Fig. 1A**). Similar results were seen for both multisensory and unisensory neurons (the neuronal-behavioral correlations remained consistently positive and negative for vestibular and visual cues, respectively). We further sorted the multisensory neurons into those with congruent and opposite vestibular and visual heading preferences (Chen et al., 2011b; Gu *et al*., 2006) with no observable differences (**Supplemental Fig. 1B**). Therefore, the contrary shifts of visual tuning in VIP seem to reflect a general feature of this cortical area, rather than an anomaly of a subgroup of neurons.

### Temporal evolution of the correlation between neuronal and behavioral shifts

The tuning curves in **Figures 3–5** were calculated using mean firing rates averaged across the stimulus duration. But the self-motion stimuli generated by the platform and optic flow followed a specific dynamical time-course, specifically, a Gaussian velocity profile and correspondingly a biphasic acceleration profile (see bottom row, **Fig. 6**). Therefore, we further examined whether the correlations between behavioral recalibration and shifts in neuronal tuning depend on the time point within the stimulus interval.

**Figure 6.**
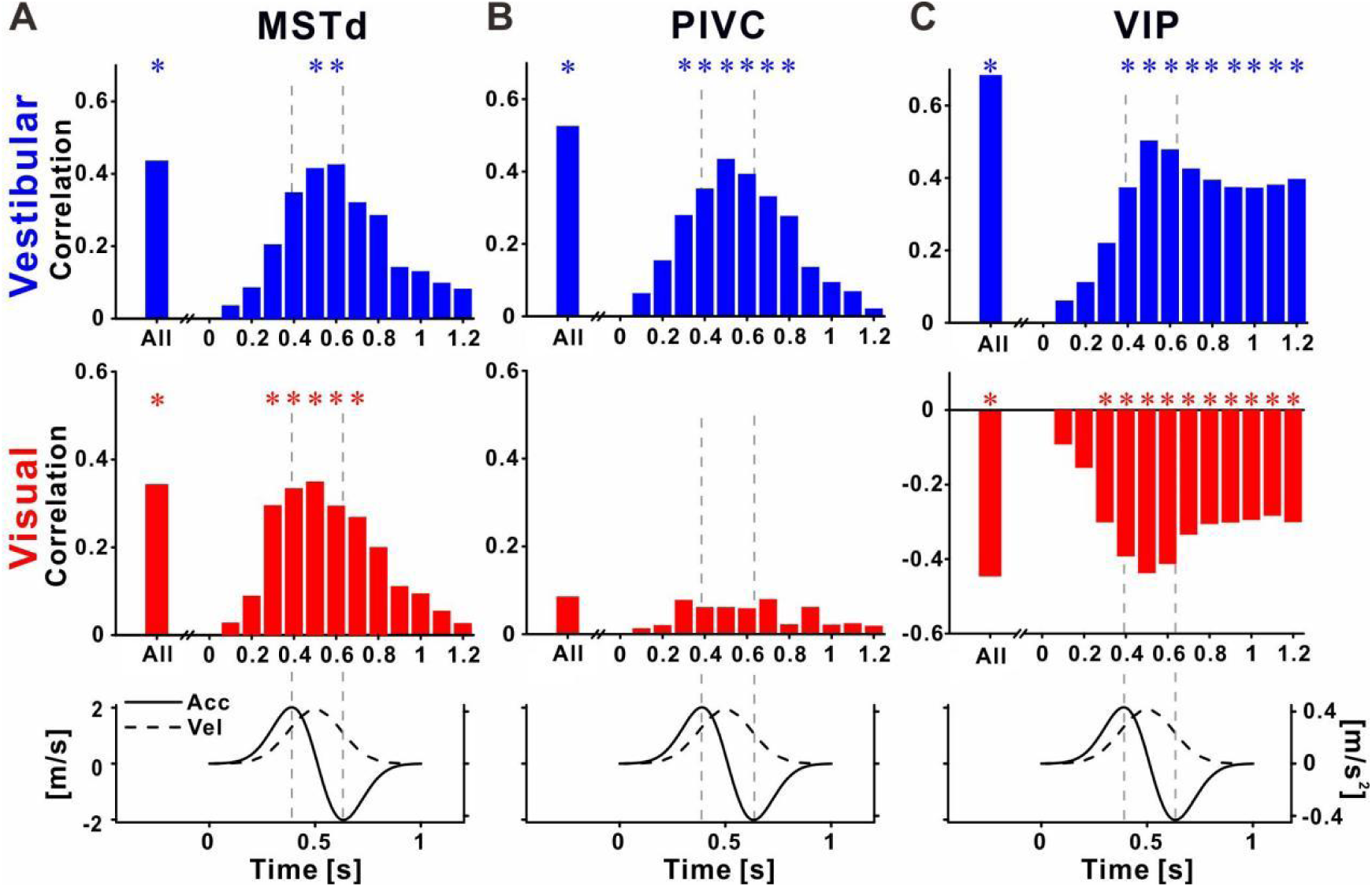
Recalibration of neuronal responses within the stimulus time-course. Correlations between neuronal and behavioral PSE shifts, using the neuronal activity at specific time-points during the stimulus, for **(A)** MSTd, **(B)** PIVC, and **(C)** VIP. Top row: vestibular (blue histograms), second row: visual (red histograms), third row: stimulus (acceleration and velocity) time-course. Vertical dashed lines mark the acceleration peaks, and ’**\***’ symbols mark significant correlations. Pearson correlation.

For MSTd neurons, positive correlations (between behavioral and neuronal shifts) were seen for both vestibular and visual cues during the stimulus (**Fig. 6A**). The profile of correlations followed the velocity profile closely. Namely correlations increased toward the middle of the stimulus, and dropped off rapidly at the end of the stimulus. Significant correlations (blue and red asterisk markers for vestibular and visual cues, respectively) were only seen around the middle of the stimulus. Thus neural recalibration in MSTd (is accordance with behavioral recalibration) is seen in velocity responses, which are transient (evident only during the stimulus).

For PIVC neurons, positive correlations (between behavioral and neuronal shifts) were seen only for vestibular cues, during the stimulus (**Fig. 6B**). Like MSTd, the correlations seemed to follow the velocity profile of the stimulus, with significant values around the middle of the stimulus (upper panel in **Fig. 6B**). Correlations in the visual condition were very weak and not significant (middle panel in **Fig. 6B**).

A very different profile was seen in VIP. Firstly, as described above, correlations between neuronal and behavioral recalibration were positive for the vestibular cue (upper panel in **Fig. 6C**) and negative for the visual cue (middle panel in **Fig. 6C**). Furthermore, the time-course of these correlations was different in VIP: they increased in size gradually (positively for vestibular and negatively for visual), reaching a maximum around the middle of the stimulus epoch (the velocity peak), but then remained elevated beyond the end of the stimulus (**Fig. 6C**). This pattern is in line with sustained neuronal activity described previously for VIP. However, here this sustained activity correlated with subsequent vestibular choices, and was contrary to visual choices. Thus the sustained activity is not generically choice related, but rather in accordance with recalibrated vestibular function.

### VIP choice signals are reduced during cross-sensory recalibration

Previous studies have found that neuronal responses in VIP are strongly influenced (perhaps even dominated) by choice signals (Chen et al., 2021; Zaidel *et al*., 2017). Hence our finding here, that neuronal tuning recalibrated contrary to behavioral shifts for the visual cue, was surprising and counterintuitive. We therefore wondered what happened to the strong choice signals for which VIP is renowned, which would predict that neuronal tuning (also for visual cues) would shift with behavior. To visualize choice tuning for an example VIP neuron, we plotted ‘choice-conditioned’ tuning curves, namely, neuronal responses as a function of heading, separately for rightward and leftward choices, (**Fig. 7**). In the pre-recalibration block vestibular responses were strongly choice related (**Fig. 7A**, left plot) – neuronal responses to the same heading stimulus were larger when followed by rightward (►, dark blue) vs. leftward (◄, cyan) choices (the dark blue line lies above the cyan line). After recalibration, the choice effect decreased (**Fig. 7A**, right plot) – the choice-conditioned tuning curves were no longer separate. Similarly, visual responses were strongly choice-related pre-recalibration, and this decreased post-recalibration (**Fig. 7B**). To quantify the choice (and sensory) components of neuronal activity, and to observe how these changed after recalibration, we applied a partial correlation analysis (Zaidel *et al*., 2017). For this example neuron, the partial choice correlation values (R_c_, presented on the plots) were reduced both for vestibular and visual cues.

**Figure 7.**
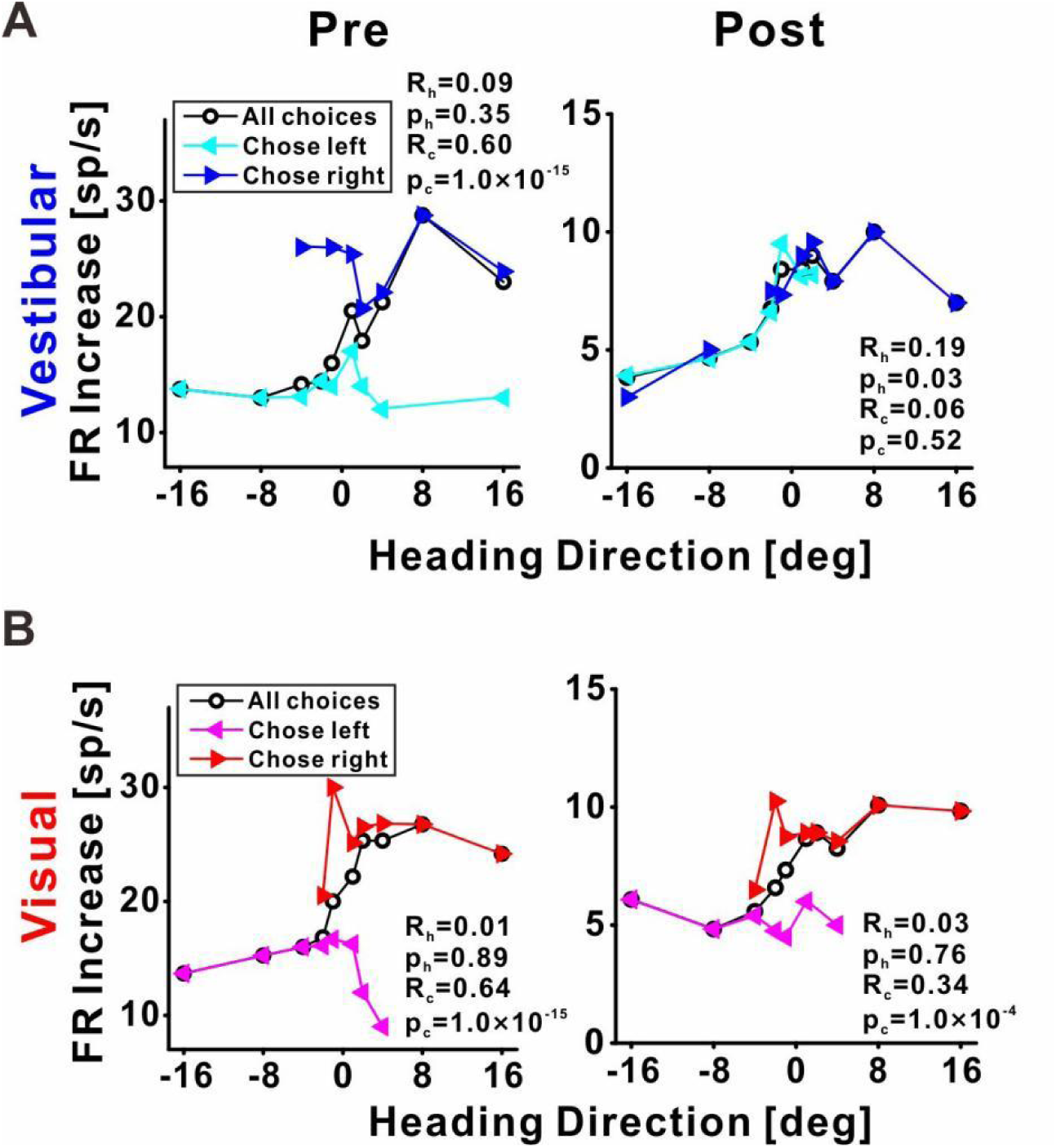
Choice tuning is reduced post-recalibration in an example VIP neuron. Neuronal responses for example VIP neuron to **(A)** vestibular and **(B)** visual heading stimuli, pre- and post-recalibration (left and right columns, respectively). Blue and cyan curves depict choice-conditioned tuning curves (neuronal responses followed by rightward and leftward choices, respectively) for the vestibular cue. Red and magenta curves depict choice-conditioned tuning curves for the visual cue. Black curves (in the corresponding plots) represent all responses (not sorted by choice). Partial heading (R_h_) and partial choice (R_c_) correlations (with corresponding p-values) are presented on the plots.

Across our sample of VIP neurons, the choice partial correlations in the post-recalibration block were significantly reduced compared to the pre-recalibration block, for both vestibular (*p* = 3.73 × 10^−3^, paired *t*-test) and visual (*p* = 4.39 × 10^−3^, paired *t*-test; **Fig. 8B**) cues. However, the heading partial correlations (R_h_) did not differ significantly from pre- to post-recalibration, neither for vestibular (*p* = 0.36, paired *t*-test) nor visual (*p* = 0.47, paired *t*-test; **Fig. 8A**) cues. For these statistical comparisons and for plotting we used the squared partial correlations (which quantify the amount of unique variance explained by choice or heading). We did not observe any changes in partial correlations in areas PIVC and MSTd (**Supplemental Fig. 2**). Lastly, there was no evidence for differences between post- and pre-recalibration baseline firing rates in any of the three areas (**Supplemental Fig. 3**; Bayes Factors (BF_10_) presented on the corresponding subplots). Thus, shifts in neuronal tuning are not explained by changes in baseline activity.

**Figure 8.**
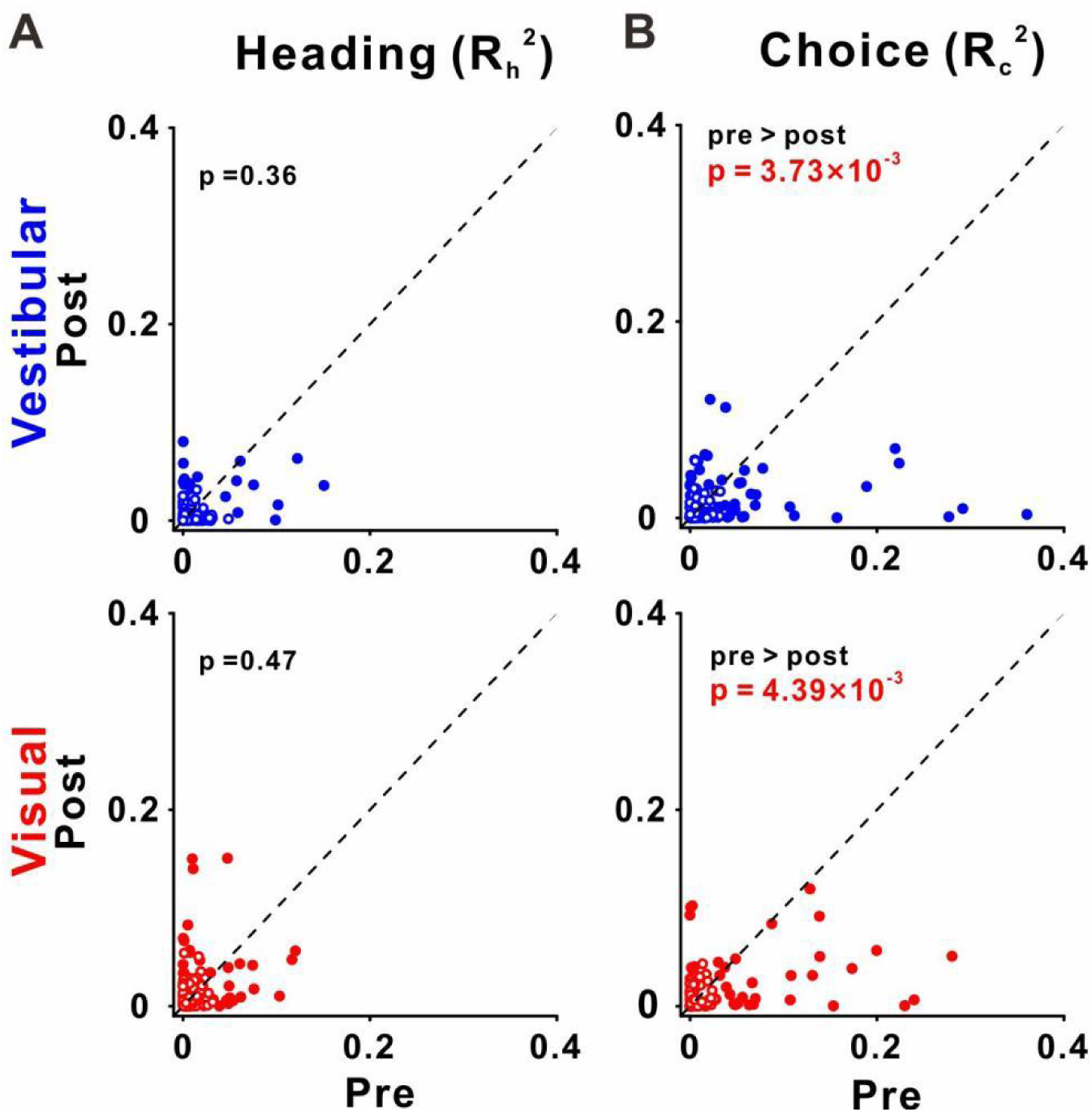
Choice tuning is reduced in VIP post-recalibration. **(A)** Heading and **(B)** choice partial correlation coefficients (squared) are depicted post- vs. pre-recalibration. Blue and red dots (top and bottom row) represent vestibular and visual cues, respectively. P-values for the hypothesis of greater pre- vs. post-recalibration values are presented on the corresponding plots. Paired t-test.

## Discussion

This study provides the first demonstration of unsupervised (cross-sensory) neuronal recalibration, in conjunction with behavioral recalibration, in single sessions. Single-neurons from MSTd, PIVC, and VIP revealed clear but different patterns of recalibration. In MSTd, neuronal responses to vestibular and visual cues recalibrated - each according to their respective cues’ perceptual shifts. In PIVC, vestibular tuning similarly recalibrated together with the corresponding vestibular perceptual shifts (the PIVC cells were not robustly tuned to visual stimuli). However, recalibration in VIP was notably different: both vestibular and visual neuronal tuning recalibrated in the direction of the vestibular perceptual shifts. Thus, visual neuronal tuning shifted, surprisingly, contrary to visual perceptual shifts. These results indicate that neuronal recalibration differs profoundly across multisensory cortical areas.

### Neural correlates of vestibular-visual recalibration

To investigate the neuronal bases of unsupervised cross-sensory recalibration, we first replicated the behavioral results from our previous study (Zaidel *et al*., 2011). Indeed, in the presence of a systematic vestibular-visual heading offset (with no external feedback) vestibular and visual cues both recalibrated in the direction required to reduce the cue conflict. And, as before, the vestibular shifts were larger compared to the visual shifts. Thus we confirmed robust recalibration of vestibular and visual cues, resulting from a systematic discrepancy between the cues’ headings in an unsupervised context (i.e., without external feedback).

Since there was no external feedback regarding which cue was (in)accurate, unsupervised recalibration is driven by the cue conflict, presumably through an internal mechanism to maintain consistency between vestibular and visual perceptual estimates (Zaidel *et al*., 2011). Accordingly, we expected to see neuronal correlates of perceptual recalibration in early multisensory areas related to self-motion perception, specifically: MSTd, which primarily responds to visual (but also vestibular) self-motion stimuli, and PIVC, which primarily responds to vestibular stimuli. We further expected that the neuronal recalibration in MSTd and PIVC would propagate to higher-level multisensory area VIP.

In MSTd, we indeed found that both visual and vestibular neuronal signals recalibrated, each in accordance with their corresponding cue’s behavioral shifts. Hence, recalibration of visual self-motion responses was observed at least at the level of MSTd, which is the primary area in the visual hierarchy to respond to large field optic flow stimuli (Britten, 2008; Britten and van Wezel, 1998; 2002; Duffy and Wurtz, 1995; Gu *et al*., 2008; Gu *et al*., 2012; Gu *et al*., 2006; Wurtz and Duffy, 1992). We cannot ascertain whether recalibration to visual responses occurred already in earlier visual regions, such as the middle temporal visual area (MT), which projects to MSTd (Maunsell and van Essen, 1983; Ungerleider and Desimone, 1986), or whether it occurred only at the level of MSTd. Because MSTd is mainly a visual area, the recalibration of vestibular signals observed in MSTd likely occurred in upstream vestibular areas that project to MSTd, such as PIVC (Chen *et al*., 2010; 2011a). Indeed, robust vestibular recalibration (that was in line with the vestibular behavioral shifts) was observed in PIVC. Hence, neuronal correlates of perceptual recalibration were observed in relatively early multisensory areas related to self-motion perception (MSTd and PIVC).

### Individualized recalibration of vestibular and visual cues

Results from this experiment exposed individualized (sensory-specific) neuronal recalibration (in MSTd and PIVC). Namely, visual and vestibular tuning curves shifted differently (in opposite directions). This provides neuronal evidence against ‘visual dominance’, even for short-term recalibration (in single sessions). Rather, it supports the idea that cross-sensory neuronal recalibration occurs also for visual (and not only for non-visual) cues.

Furthermore, sensory-specific recalibration of visual and vestibular tuning implies that the brain has mechanisms to separately monitor and recalibrate individual cues. Cue-specific shifts in neuronal tuning were not seen during supervised recalibration, because the cues largely shift together, in response to external feedback (Zaidel *et al*., 2021). Even though in the supervised condition both unsupervised and supervised shifts are in operation (superimposed, Zaidel et al. 2013), the supervised (yoked) component is large and dominates, thereby obscuring the individualized (unsupervised) shifts. Here, without external feedback, we were able to detect individualized shifts of the different cues, not previously observed. This exposes neuronal mechanisms to maintain internal consistency between vestibular and visual cues. This dynamic cross-sensory plasticity may underlie our adept ability to adapt to sensory conflict commonly experienced in many modes of transport (on land, at sea, or in flight).

### Contrary recalibration in higher-level area VIP

VIP is a higher-level multisensory area (Bremmer et al., 2002; Colby et al., 1993; Duhamel et al., 1998; Schlack et al., 2002; Schlack et al., 2005; Schroeder and Foxe, 2002) with clear vestibular and visual heading selectivity (Chen *et al*., 2011a; b). But the nature of these self-motion signals in VIP is not fully understood. In contrast to our prediction that recalibrated signals in MSTd and PIVC would simply propagate to VIP, we found a different and unexpected pattern of recalibration in VIP. While vestibular tuning recalibrated in line with vestibular perceptual shifts (like MSTd and PIVC), visual tuning recalibrated opposite in direction to the visual perceptual shifts (and opposite in direction to MSTd visual recalibration). These findings indicate that visual responses in VIP do not reflect a simple feed-forward projection from MSTd. They also suggest that visual responses in VIP are not decoded for heading perception (otherwise these would not recalibrate in opposite directions). This interpretation is in line with findings that inactivation (Chen *et al*., 2016) and microstimulation (Yu and Gu, 2018) in VIP do not affect perceptual decisions. Thus, the convergence of visual and vestibular signals in VIP likely serves purposes other than cue integration.

The results here also shed new light on the neuronal shifts observed in VIP during supervised recalibration (Zaidel *et al*., 2017). There, because behavioral responses shifted in the same direction for both cues, it was reasonable to interpret visual and vestibular tuning shifts in accordance with their corresponding cue shifts. However, the results here indicate that yoking of visual and vestibular tuning is observed in VIP irrespective of the paradigm (supervised or unsupervised). Hence yoked recalibration is a feature of VIP, not just supervised recalibration.

We previously found strong choice-related activity in VIP neurons (Zaidel *et al*., 2017). Accordingly, we considered that shifts in VIP neuronal tuning (during supervised calibration) might simply reflect the altered choices (Zaidel *et al*., 2021). However, choice-related activity cannot explain the results here, because the predicted shifts in neuronal tuning would be in the same direction as the altered choices (behavioral shifts), whereas we found contrary visual recalibration. To understand contrary shifts that could arise despite strong choice-related activity in VIP, we investigated choice tuning in VIP neurons. We found that choice tuning in VIP decreased during unsupervised calibration. This allowed contrary shifts to be exposed. It also opens up new and fascinating questions regarding the purpose of contrary visual recalibration in VIP.

Because visual and vestibular tuning in VIP both shifted in the same direction (in accordance with vestibular behavioral shifts) we speculate that VIP recalibration reflects a global shift in the reference frame, following vestibular recalibration. This notion is consistent with suggestions that VIP encodes self-motion in head or body-centered coordinates (Chen et al., 2013b; 2018; Zhang et al., 2004). Accordingly, visual responses in VIP are transformed into a vestibular-recalibrated space. This leads to a remarkable dissociation between visual tuning in VIP and MSTd. Interestingly, visual self-motion perception follows the MSTd (not VIP) recalibration. This is in line with a causal connection between MSTd and visual heading discrimination (Britten and van Wezel, 1998; Gu *et al*., 2012). What purpose might such visual signals in VIP serve? One possible idea is that they might reflect an expectation signal – e.g., predicted vestibular or somatosensory sensation, based on the current visual signal. During combined stimuli (in the recalibration and post-calibration blocks), the visual signal always appeared together with the vestibular sensory input. Thus, if visual responses in VIP reflect vestibular expectations, then these would shift together with vestibular (rather than visual) recalibration.

### Limitations and future directions

Our results revealed correlations between neuronal recalibration and cross-sensory behavioral recalibration. However, they do not implicate any causal connections. Therefore, whether these cortical areas are actively involved in cross-sensory recalibration, vs. simply reflecting the recalibrated signals, requires further research. To probe more directly for causal links, direct manipulation of neuronal activity might be required. For example, would reversible inactivation or microstimulation (of one or a combination of these multisensory areas) eliminate (or bias) unsupervised recalibration? In addition, future studies are needed to examine how the systematic error between vestibular and visual heading signals is detected. This likely involves additional brain areas, for example, the cerebellum, implicated in internal-model-based error monitoring (Markov et al., 2021; Rondi-Reig et al., 2014), and/or the Anterior Cingulate Cortex (ACC), implicated in conflict monitoring (Bush et al., 2000; Holroyd and Coles, 2002). Thus, a wide-ranging effort to record and manipulate neural activity across a variety of brain regions will be necessary to tease apart the circuitry underlying this complex and important function.

The most surprising and intriguing finding in this study was the contrary recalibration of visual tuning in VIP. We propose that yoked recalibration of visual and vestibular responses in VIP (despite differential behavioral recalibration) might reflect a global shift in vestibular space. Accordingly, we suggest that visual responses in VIP might reflect an expectation signal (in vestibular space), e.g., a simulation of the expected corresponding vestibular response (or integrated position, because VIP responses are sustained beyond the stimulus period). However, this idea is speculative, and the data from this study cannot address this question. Hence, further research is needed to investigate this idea, for example, by conditioning expectations for vestibular motion on other (non-motion) cues, and investigating whether these cues can induce simulated vestibular responses. If this hypothesis turns out to be true, it could greatly contribute to our understanding regarding the functions of the parietal cortex, and the brain mechanisms of perceptual inference.

### Concluding remarks

This study exposed individualized (sensory-specific) recalibration of neuronal signals, resulting from a cross-sensory (visual-vestibular) cue conflict. It further revealed profound differences in neuronal recalibration across multisensory cortical areas MSTd, PIVC and VIP. The results therefore provide novel insights into adult multisensory plasticity, and deepen our understanding regarding the different functions of these multisensory cortical areas.

## Methods

### Subjects and surgery

Three male rhesus monkeys (*Macaca mulatta*, monkeys D, B, and K) weighing 8–10 kg participated in the experiment. Monkeys were first trained to sit in a custom primate chair and gradually exposed to the laboratory environment. Then they chronically implanted a head-restraint cap and a sclera coil for measuring eye movement. After full recovery, monkeys were trained to perform experimental tasks. All animal surgeries and experimental procedures were approved by the Institutional Animal Care and Use Committee at East China Normal University (IACUC protocol number: Mo20200101).

### Equipment setup and motion stimuli

During the experiments, monkeys were head-fixed and seated in a primate chair which was secured to a six-degree of freedom motion platform (Moog, East Aurora, NY, USA; MB-E-6DOF/12/1000Kg). The chair was also inside the magnetic field coil frame (Crist Instrument Co., Inc., Hagerstown, MD, USA) mounted on the platform for measuring eye movement with the sclera coil technique (for details, see Zhao et al., 2021).

Vestibular stimuli were delivered by the motion platform (for details, see Gu et al., 2006; Chen et al., 2013; Zhao et al., 2021). Visual stimuli were presented on a large computer screen (Philips BDL4225E, Royal Philips, Amsterdam, Netherlands), attached to the field coil frame. The display (62.5 cm × 51.5 cm) was viewed from a distance of 43 cm, thus subtending a visual angle of 72° × 62°. The sides of the coil frame were covered with a black enclosure, so the monkey could only see the visual stimuli on the screen (Gu *et al*., 2006; Zhao et al., 2021). The display had a pixel resolution of 1920 x 1080 and was updated at 60 Hz. Visual stimuli were programmed in OpenGL to simulate self-motion through a 3D cloud of “stars” that occupied a virtual cube space 80 cm wide, 80 cm tall, and 80 cm deep centered on the central point on the screen. The random-dot density was 0.01/cm^3^ (each “star” comprised a triangle with base by height: 0.15 cm × 0.15 cm). Monkeys wore custom stereo glasses made from Wratten filters (red #29 and green #61; Barrington, NJ, USA), such that the optic flow stimuli could be rendered in three dimensions as red-green anaglyphs.

The self-motion stimulus was either vestibular-only, visual-only, or combined (visual and vestibular stimuli). In the vestibular-only condition, there was no optical flow on the screen and the monkey was translated by the motion platform. In the visual-only condition, the motion platform remained stationary while the optic flow was presented on the screen. For the combined condition, the monkey experienced both translation motion and optic flow simultaneously. Each stimulus motion followed a Gaussian profile with a duration of 1 s, and an amplitude of 13 cm (bottom row, **Fig. 6**). The peak velocity was 0.41 m/s, and the peak acceleration was 2.0 m/s^2^.

### Task and recalibration protocol

The monkeys were trained to report their direction of translation with a two-alternative forced-choice (2AFC) heading discrimination task (for details, see Gu et al., 2008; Chen et al., 2013). In each trial, the monkey experienced a primarily forward motion with a small leftward or rightward component. During stimulation, the animal was required to maintain fixation on a central point within a 3° × 3° window. At the end of the trial (after a 300ms delay period beyond the end of the stimulus), the monkeys needed to make a saccade to one of two choice targets (located 5° to the left and right of the central fixation point) to report their perceived motion as leftward or rightward relative to straight ahead. The saccade endpoint had to remain within 2.5° of the target for at least 150 ms to be considered a valid choice. Correct responses were rewarded with a drop of liquid.

To elicit recalibration, we used a similar unsupervised cue-conflict recalibration protocol previously tested behaviorally in humans and monkeys (Zaidel *et al*., 2011). Each experimental session consisted of three consecutive blocks, as described here below.

#### Pre-recalibration block

This block was used to deduce the baseline performance (psychometric curve) of each modality for the monkeys, thus only a single cue (vestibular-only or visual-only) stimulus was presented (**Fig. 1A**). Across trials, the heading angle was varied in small steps around straight ahead. Ten logarithmically spaced heading angles were tested for each monkey (±16°, ±8°, ±4°, ±2°, and ±1°). To get monkeys accustomed to not getting a reward for all the trials, we rewarded the monkeys with a 95% probability for correct choices and didn’t reward them for incorrect choices.

#### Recalibration block

Only combined vestibular-visual cues were presented in this block (**Fig. 1B**). There was a discrepancy (Δ) between the vestibular and visual cues, which was introduced gradually from ± 2° to ± 10° with steps of 2°, and then held at ± 10° for the rest of the block. This gradual introduction was designed to prevent monkeys from realizing the discrepancy. The sign of Δ represents the orientation of discrepancy: positive Δ (i.e. Δ^+^) indicates vestibular cue to the right and visual cue to the left, and vice versa for negative Δ (i.e. Δ^−^). Every session used only one sign, positive or negative. The combined cue heading was defined as the midpoint between the vestibular and visual headings, such that each (vestibular/visual) heading was offset to the right and left (or left and right) in relation to the combined heading. The same ten heading angles as in the pre-recalibration block were used. Unlike the pre-recalibration block, monkeys only needed to maintain fixation on the central point during the stimulus presentation and didn’t need to make choices at the end of trials. They were rewarded for all the trials for which they maintained fixation. 7∼10 repetitions were run for each Δ increment, and an additional 10∼16 repetitions were run for maximum Δ (±10°).

#### Post-recalibration block

During this block, performance of the individual (visual/vestibular) modalities was once again tested using single modality trials (as in the pre-recalibration block). Responses to these trials were used to measure recalibration. The single cue trials were interleaved with combined-cue trials (with a 10° discrepancy, like the end of the recalibration block, **Fig. 1C**). The combined cue trials were interleaved to maintain the recalibration while it was measured (for details, see Zaidel et al., 2011). To avoid perturbing the recalibrated behavior, we adjusted the reward probability for single-cue trials as follows: if the single cue heading was of relatively large magnitude, such that, if it were part of a combined cue trial also the other cue would lie to the same side (right or left), monkeys were rewarded as in the pre-recalibration block (95% probability reward for correct choices; no reward for incorrect choices). If, however, the heading for other modality would have been to the opposite side, the monkeys were rewarded stochastically (70% reward probability, regardless of their choices).

### Electrophysiological recordings

We recorded extracellular activity from isolated single neurons in areas MSTd, PIVC, and VIP using tungsten microelectrodes (Frederick Haer Company, Bowdoin, ME, USA; tip diameter ∼3 *μ*m; impedance, 1∼2 MΩ at 1 kHz). The microelectrode was advanced into the cortex through a transdural guide tube, using a hydraulic microdrive (Frederick Haer Company). Raw neural signals were amplified, band-pass filtered (400–5000 Hz), and digitized at 25 kHz using the AlphaOmega system (AlphaOmega Instruments, Nazareth Illit, Israel). The spike times sorted online along with all behavioral events were collected with 1 ms resolution using the Tempo system for offline analysis. If the online sorting was not adequate, offline spike sorting was performed.

The target areas (VIP, PIVC, and MSTd) were identified based on the patterns of gray and white matter transitions, magnetic resonance imaging (MRI) scans, stereotaxic coordinates, and physiological response properties as described previously (MSTd: Gu et al., 2006; PIVC: Chen et al., 2010; VIP: Chen et al., 2011).

## Data analysis

Data analysis was performed with custom scripts in Matlab R2016a (The MathWorks, Natick, MA, USA). Psychometric function plots were constructed by plotting the proportion of “rightward” choices as a function of heading angle and then fitted with a cumulative Gaussian distribution function using the *psignifit* toolbox for MATLAB (version 2.5.6). For each experimental session, separate psychometric functions were constructed for visual and vestibular conditions before and after recalibration. The psychophysical threshold and point of subjective equality (PSE) were defined as the SD (σ) and mean (μ), respectively, deduced from the best-fitting function. The PSE represents the heading angle of equal right/left choice proportion, i.e., perceived straight ahead, also known as the bias. The vestibular/visual recalibration effect was calculated for each session by subtracting the PSE value of the pre-recalibration from that of the post-recalibration PSE.

Neuronal heading tuning curves were constructed (pre/post recalibration block and vestibular/visual cue) by computing the average FR (in units of spikes/s, the baseline FRs subtracted) for each heading over the stimulus presentation (t=0-1s). A neuron was considered tuned to vestibular or visual cue if the linear regression of FR vs. heading (over the narrow range −16° to 16°) had a significant slope (p < 0.05, Pearson’s correlation). When calculating the group effects of recalibration for a vestibular or visual cue, we only considered cells with significant tuning either pre- or post-recalibration. This resulted in 49 and 66 (of 118 recorded) VIP neurons tuned to vestibular and visual cues, respectively (31 of which were tuned to both); 60 (of 160 recorded) PIVC neurons tuned to vestibular cues; 23 and 65 (of 83 recorded) MSTd neurons tuned to vestibular and visual cues, respectively.

To estimate neural recalibration (for comparison to behavioral recalibration) we constructed neurometric functions (Chen *et al*., 2013a; Fetsch *et al*., 2012; Gu *et al*., 2008; Gu *et al*., 2007) for the pre-recalibration and post-recalibration data (each calculated after subtracting the mean baseline firing rate respectively). Specifically, both the pre-recalibration and post-recalibration data were normalized (z-scored) by subtracting the pre-recalibration mean response and dividing by the pre-recalibration SD across stimulus repetitions. Then ROC (receiver operating characteristic) analysis was used to compute the ability of an ideal observer to discriminate between the z-scored responses (for each heading) and 0 ° (straight ahead). These ROC values were fitted with a cumulative Gaussian function (like for behavioral psychometrics), and neuronal recalibration was measured by the difference in PSE (as done for behavior).

To assess neuronal recalibration at different time points during the stimulus, we calculated response metrics in 200 ms time windows, starting at stimulus onset, and shifted in steps of 100 ms. The time index (the center of the window) ranged from t = 0.1 s to t = 1.2 s (relative to stimulus onset). This range did not include the saccade, which could only take place after t = 1.3 s because of the delay period (300ms) that was at the end of the stimulus.

### Partial correlation analysis

To disassociate the unique contributions of heading stimuli and choices to the neural responses (FRs), we computed Pearson partial correlations between these variables (for details, see Zaidel et al., 2017; Chen et al., 2021). This produced a heading partial correlation, R_h_, that captured the linear relationship between firing rate (FR) and heading (H) given the monkey’s choice (C), as well as a choice partial correlation, R_c,_ that captured the relationship between firing rate and choice given the stimulus heading. Partial correlations were calculated over the entire 1 s stimulus duration. Positive heading partial correlations indicate that firing rates were greater for rightward than leftward headings (given the choices). Likewise, positive choice partial correlations indicate that firing rates were greater for choices made to the right than choices made to the left (given the stimulus headings).

### Statistical Analysis

To evaluate differences in monkeys’ behavior (PSE), heading, or choice partial correlations, between pre- and post-recalibration, we used paired t-tests. Possible differences in spontaneous (baseline) firing rates between pre- and post-recalibration were evaluated using Bayesian paired-samples t-tests (BF_10_ values). Statistical analysis was conducted using the open-source statistical software program JASP (Version 0.16.3).

## Acknowledgments

This work was supported by grants from the “technology innovation 2030--major projects” on brain science and brain-like computing of the Ministry of Science and Technology of China (No. 2021ZD0202600), the National Basic Research Program of China (No. 32171034) to A.C., and the ISF-NSFC joint research program to A.C. (No. 32061143003) and A.Z. (No. 3318/20). We thank Prof. Dora Angelaki for the helpful comments. We are also grateful to Minhu Chen for outstanding computer programming.

## Competing interests

Authors declare no competing interests.

## Data and Code availability statement

The data and analysis code for this study have been uploaded to github and can be found at https://github.com/FuZengBio/Recalibration.

## Additional files

Supplementary files

## Supplementary Material

**Supplemental Figure 1.**
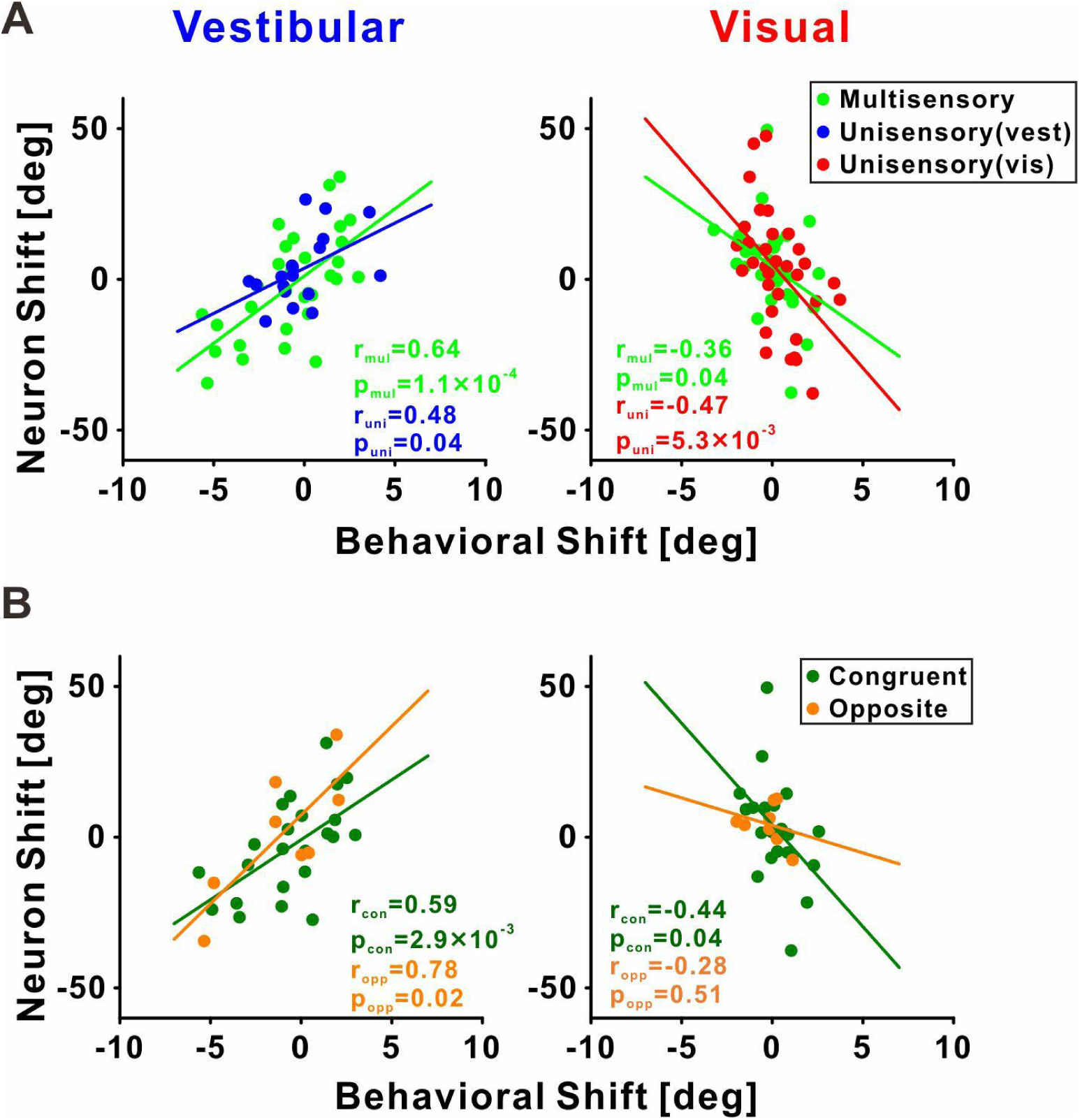
Neuronal vs. behavioral shifts by neuron type in area VIP. (**A)** Neurons with multisensory (green) and unisensory (blue and red, for vestibular and visual, respectively) tuning. (**B)** Multisensory neurons with congruent, or opposite, vestibular and visual tuning. The neuronal shifts were positively correlated with the behavioral shifts for the vestibular cue (left column), and negatively correlated with the behavioral shifts for the visual cue (right column). Pearson correlation coefficients are presented on the corresponding plots.

**Supplemental Figure 2.**
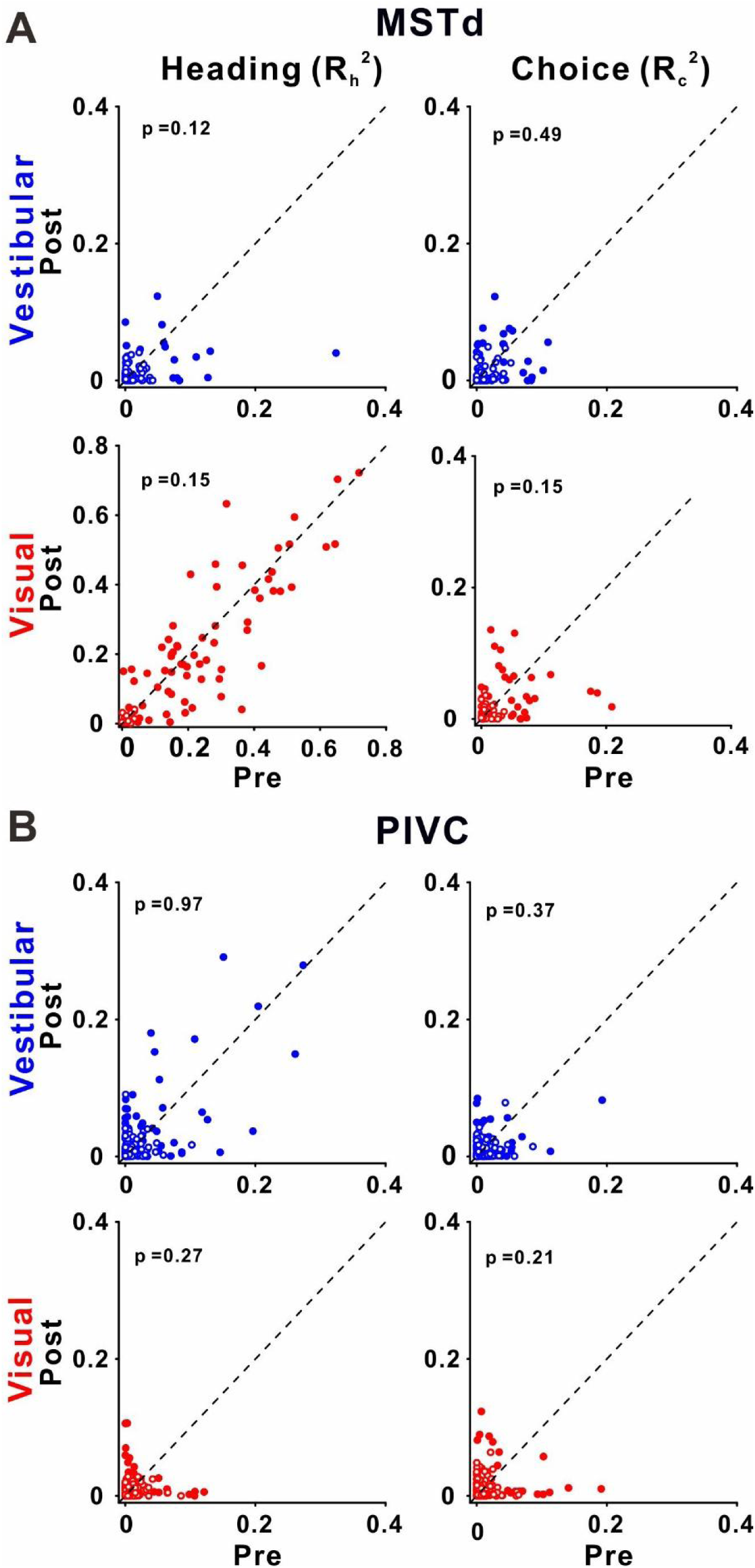
Choice and heading partial correlations in areas MSTd and PIVC. Plotting conventions are the same as in Figure 8.

**Supplemental Figure 3.**
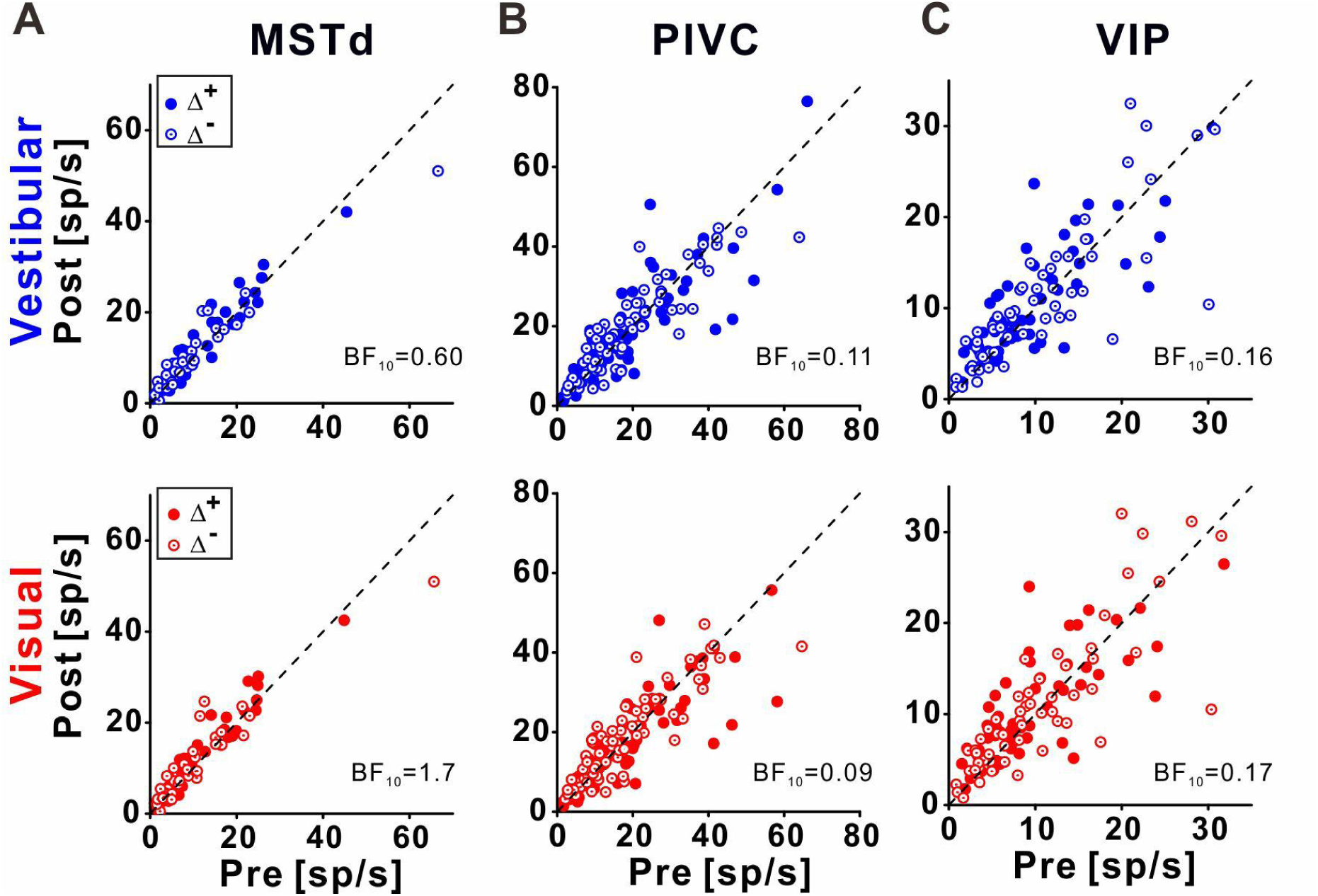
Baseline firing rates in areas MSTd, PIVC and VIP. The baseline firing rates post- vs. pre-recalibration are plotted for vestibular (upper panel) and visual (bottom panel) cues. Solid symbols represent Δ^+^ and open symbols represent Δ^−^. Bayes factors (BF_10_) < ⅓ (as for PIVC and VIP) provide substantial evidence against a change in baseline firing rates. Bayes factors between ⅓ and 3 (as for MSTd) are inconclusive (provide no substantial evidence for, or against, changes).

## Notes

### Competing Interest Statement

The authors have declared no competing interest.

